# Nucleated transcriptional condensates amplify gene expression

**DOI:** 10.1101/737387

**Authors:** Ming-Tzo Wei, Yi-Che Chang, Shunsuke F. Shimobayashi, Yongdae Shin, Clifford P. Brangwynne

## Abstract

Liquid-liquid phase separation is thought to underly gene transcription, through the condensation of the large-scale nucleolus, or in smaller assemblies known as transcriptional hubs or condensates. However, phase separation has not yet been directly linked with transcriptional output, and our biophysical understanding of transcription dynamics is poor. Here, we utilize an optogenetic approach to control condensation of key FET-family transcriptional regulators, particularly TAF15. We show that amino acid sequence-dependent phase separation of TAF15 is enhanced significantly due to strong nuclear interactions with the C-terminal domain (CTD) of RNA Pol II. Nascent CTD clusters at primed genomic loci lower the energetic barrier for nucleation of TAF15 condensates, which in turn further recruit RNA Pol II to drive transcriptional output. These results suggest a model in which positive feedback between key transcriptional components drives intermittent dynamics of localized phase separation, to amplify gene expression.

---

Cells organize complex biochemical reactions through compartmentalization into various organelles. Many of these organelles are membrane-less condensates, which form through liquid-liquid phase separation (LLPS), in which biomolecules condense into dynamic liquid droplets ^1–4^. Nuclear condensates are thought to play key roles in regulating the flow of genetic information and include a diverse set of structures ranging from large scale assemblies such as nuclear speckles and the nucleolus, to structures that form on smaller length scales such as Cajal bodies, promyelocytic leukemia (PML) bodies, and gems.

The concept that LLPS can drive transcriptional activity has recently received intense attention. The nucleolus is a large nuclear condensate which assembles around actively transcribing ribosomal RNA (rRNA) loci ^5, 6^, and is thought to facilitate the processing and maturation of rRNA into pre-ribosomal particles. Smaller transcriptional condensates have been proposed to assemble throughout the genome to facilitate transcription of non-ribosomal genes through the RNA polymerase II (Pol II) machinery ^7–10^. These nanoscale transcriptional condensates have been hypothesized to facilitate enhancer-promoter interactions ^11–13^, possibly through the generation of localized force through surface-tension driven droplet coalescence ^14^.

Transcriptional condensates and other nuclear bodies are enriched in nucleic acid-binding proteins containing large intrinsically disordered proteins or regions (IDPs/IDRs), closely related to low complexity sequences (LCS) and prion-like domains (PrLD). The amino acid sequences of IDRs encode an intrinsic preference for conformational heterogeneity and do not fold into a single three-dimensional structure. IDRs have been shown in numerous studies to promote phase separation ^15–19^, although how specificity emerges from their sequence-encoded interactions is still largely unclear.

Of particular interest are members of the FET protein family, consisting of the proteins FUS (fused in sarcoma or translocated in liposarcoma), EWS (Ewing’s sarcoma), and TAF15 (TATA-binding protein-associated factor 2N), which play roles not only in transcription but also in pre-mRNA splicing, DNA repair, and mRNA transport in neurons ^20^. FUS, EWS, and TAF15 all contain an N-terminal IDR enriched in uncharged polar amino acids and aromatic amino acids (Fig. S1), an RNA-binding motif (RNA-recognition motif, RRM), and a R/G (arginine/glycine)-rich domain (Fig. 1A). While recent *in vitro* studies have begun to explore the role of these amino acids in defining molecular interactions and specificity ^21–23^, this remains poorly understood, particularly within living cells. Moreover, whether and how sequence-encoded phase behavior can determine functional transcriptional outputs, and their reported intermittent dynamics ^24^, remains to be tested.

**Fig. 1.**
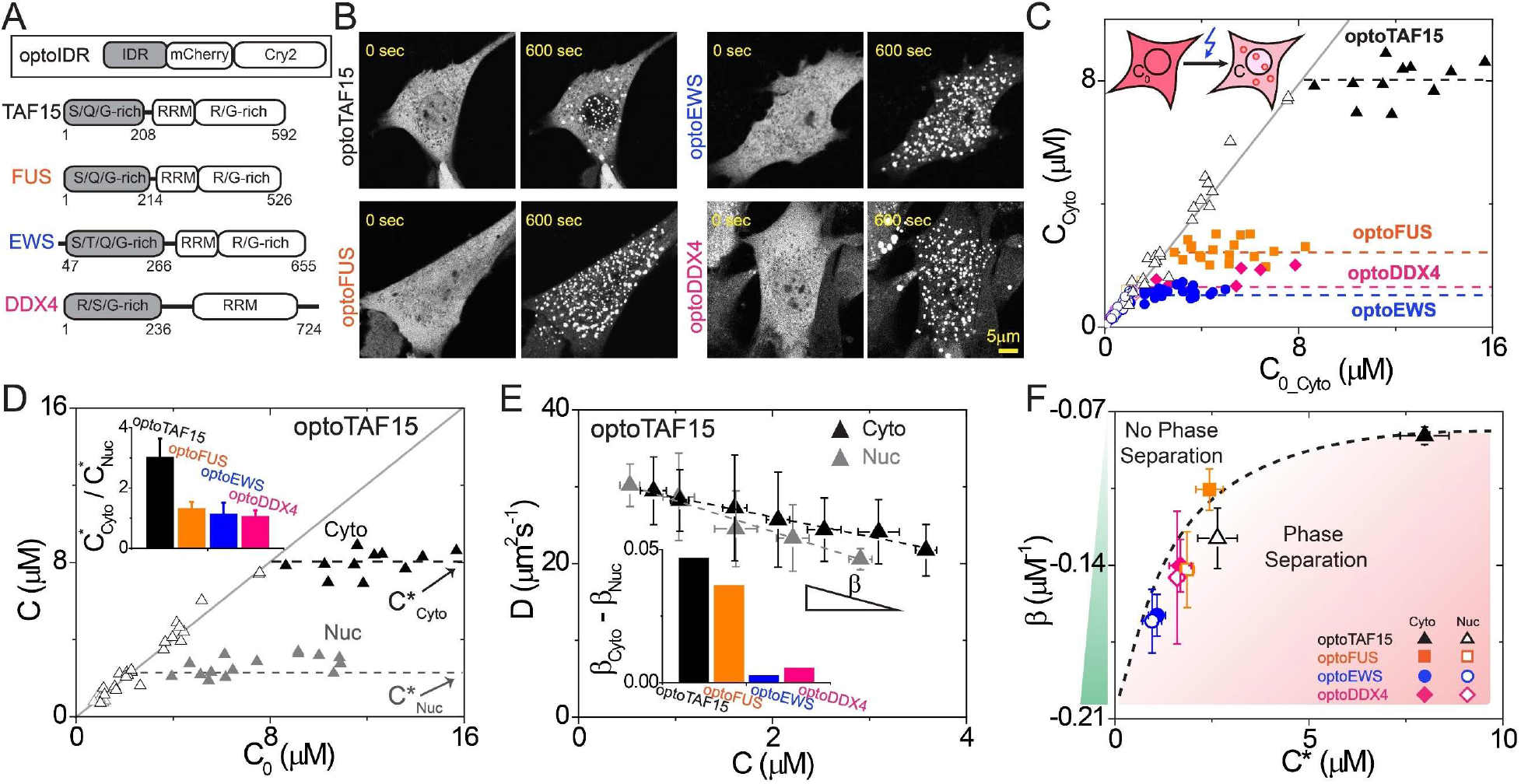
Molecular interaction strength drives intracellular phase behavior. (**A**) Schematic diagram of optogenetic platform. Each “optoIDR” construct consists of an N-terminal IDR (from TAF15, FUS, EWS, and DDX4) fused to fluorescent protein mCherry and the Cry2 domain. (**B**) Blue light illumination leads to induction of optoIDR condensates. Time since the start of illumination is shown with 0 sec indicating just before illumination. Scale bar is same for all images. (**C**) Steady-state cytoplasmic optoIDR concentration outside clusters (C_Cyto_) from individual cells under blue light illumination as a function of optoIDR initial concentration (C_0_Cyto_; i.e., cytoplasmic concentration measured before exposure to blue light). Solid and open symbols represent individual cells with or without light-activated assemblies respectively; solid line has slope of 1 and y-intercept of 0. (**D**) Similar plot as (C) for cytoplasmic (black) and nuclear (gray) optoTAF15, which exhibits a higher saturation concentration in cytoplasm (C*_Cyto_) than nucleoplasm (C*_Nuc_). (Inset) Fold-change of cytoplasmic to nucleoplasmic saturation concentration of each optoIDR construct tested. Error bars, errors propagated from standard deviations. (**E**) Diffusion coefficient of optoIDR (FCS measurements in dark) as a function of the protein concentration. Data are plotted as mean ± standard deviation (n = 5-10 cells). The dashed lines are least-square fits of optoTAF15 data to *Eqn*. 1. (Inset) OptoTAF15 exhibits the greatest difference of molecular interaction strengths between the cytoplasm (β_Cyto_) and nucleoplasm (β_Nuc_). (**F**) Intracellular phase diagram generated by plotting IDR molecular interaction strength against the optoIDR saturation concentration (C*). Horizontal error bars, standard deviations; vertical error bars, standard errors from fitting. The dashed curve is drawn qualitatively.

## Results

### Molecular interaction strength drives intracellular phase behavior

The valence of IDR-rich proteins is often enhanced through binding to multiple genomic elements ^17, 18^, or through direct oligomerization ^25^. To address the question of the sequence-dependent transcriptional function of FET family proteins, we took advantage of the recently developed optoDroplet system ^26^, which utilizes blue-light controlled oligomerization of the Cryptochrome 2 (Cry2) protein to control IDR oligomerization, and thus drive phase separation ^20, 27^. In agreement with earlier work, we find that blue light drives phase separation of an optoFUS construct into liquid-like droplets in both the cytoplasm and nucleoplasm of NIH 3T3 cells (Fig. 1B and Fig. S2). Consistent with the physics of classical phase separation, blue light-illuminated cells expressing different optoFUS concentrations reveal that the cytoplasmic concentration (C_cyto_) begins to plateau at a saturation concentration (C^*^) at roughly C*_Cyto_= 2.4 ± 0.4 μM (Fig. 1C). We observe similar behavior with an optoEWS construct and an optoTAF15 construct, which consist of Cry2 fused to the IDR/LCS regions of EWS and TAF15, respectively (Fig. 1C). Interestingly, however, optoTAF15 only phase separates in the cytoplasm at significantly higher concentrations, C*_Cyto_ = 8.0 ± 0.6 μM, which is roughly 3-fold higher than the saturation concentration for nucleoplasmic optoTAF15 (2.6 ± 0.5 μM; Fig. 1D). Thus, nucleoplasmic optoTAF15 has a significantly enhanced tendency to phase separate in the nucleoplasm compared to the cytoplasm, whereas other constructs tested exhibit only moderate differences between the cytoplasmic and nucleoplasmic saturation concentration (Fig. 1D, inset).

To examine the biophysical origins of this surprising TAF15 phase behavior, we utilize fluorescence correlation spectroscopy (FCS) to estimate molecular interaction strength. An effective interaction strength, β, is determined by measuring protein diffusivity as a function of protein concentration, and analyzing the data using the relation D = D_0_ (1 + βC) (*Eqn*. 1) where D is protein diffusivity, D_0_ is the diffusivity at infinite dilution, and C is the protein concentration ^28, 29^. In the absence of blue light illumination, optoTAF15 diffusivity decreases with increasing protein concentration (Fig. 1E), which indicates an attractive molecular interaction. However, optoTAF15 exhibits significantly stronger interactions in the nucleoplasm compared to the cytoplasm (i.e. β_Nuc_ is more negative than β_Cyto_). OptoFUS also has slightly stronger interactions in the nucleoplasm, but not nearly as large as for optoTAF15. OptoEWS and a control construct optoDDX4 both exhibit no significant difference between their cytoplasmic and nucleoplasmic interaction strengths (Fig. 1E inset). For all constructs tested, the apparent molecular interaction strength in the absence of blue light illumination can be plotted against the saturation concentration for optoIDRs under blue light illumination, yielding a common intracellular phase boundary (Fig. 1F). These findings suggest that IDR-driven phase separation exhibits remarkably universal features, and that the enhanced tendency for optoTAF15 to phase separate in the nucleoplasm compared with the cytoplasm reflects its correspondingly stronger intermolecular interactions there (Fig. 1E and 1F).

### Transcriptional condensates colocalize with and recruit unphosphorylated CTD

We next sought to understand the origin of TAF15’s increased interaction strength and preferential condensation in the nucleoplasm. TAF15 is a key transcriptional regulator and has been shown to interact with the C-terminal domain (CTD) of Pol II ^27, 30^. We co-expressed the optoIDR constructs along with EGFP-tagged Pol II CTD construct consisting of a 52-repeat of the heptad consensus amino acid sequence YSPTSPS. We find that optoTAF15 droplets exhibit a nearly 4-fold preferential partitioning of CTD (Fig. 2A and 2B). As a negative control, there is no preferential partitioning of EGFP-CTD into induced optoDDX4 droplets. OptoEWS droplets similarly exhibit no CTD recruitment, while optoFUS exhibits a 2-fold enrichment of CTD (Fig. 2A and 2B).

**Fig. 2.**
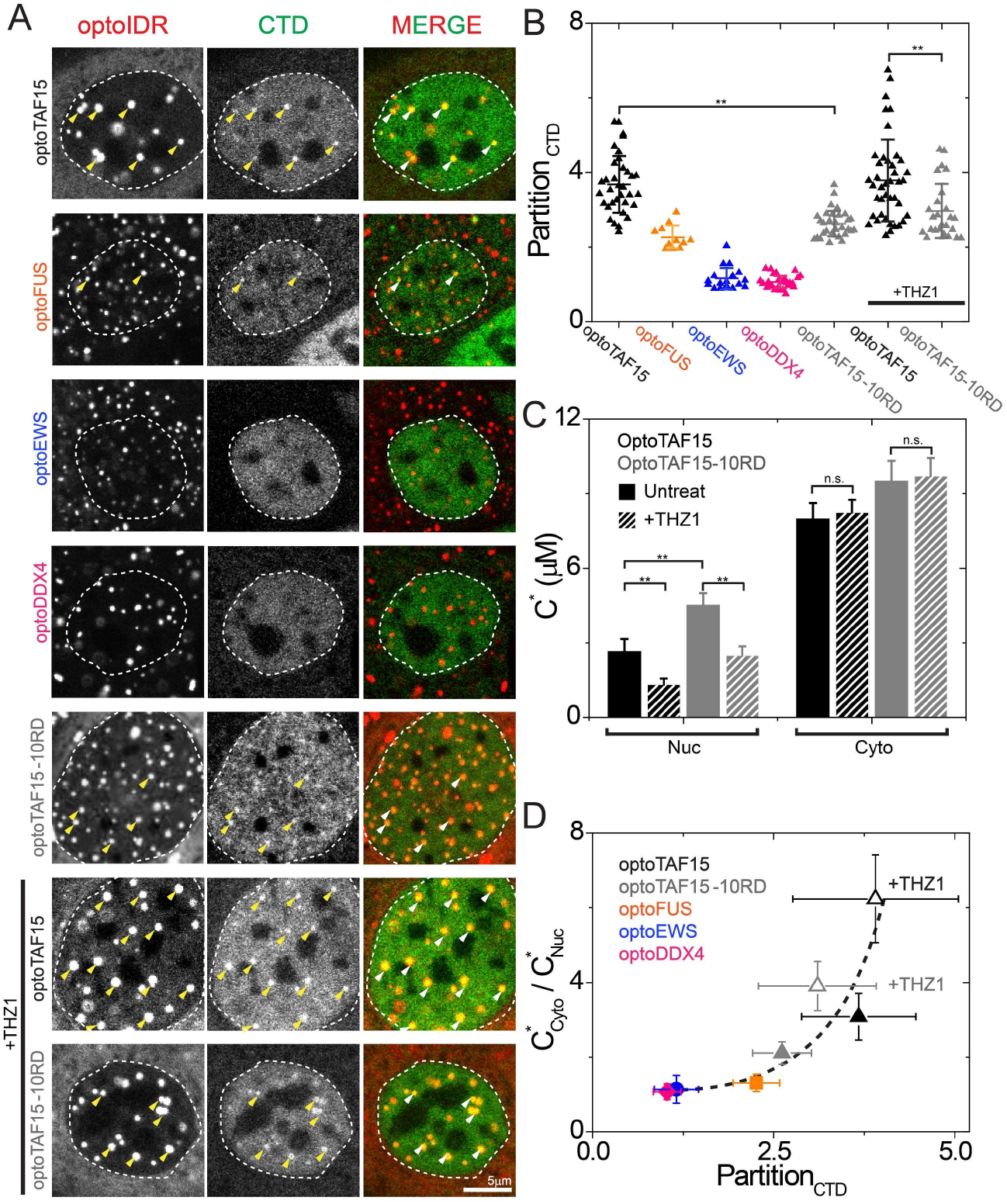
Transcriptional condensates colocalize and recruit unphosphorylated CTD. (**A**) Fluorescent images of live cells expressing optoIDRs and EGFP-CTD after blue-light illumination. Cell nucleus is outlined by dotted line. Arrowheads point to optoIDR droplets colocalized with recruit CTD. Scale bar same for all images. (**B**) CTD partition coefficients in various optoIDR droplets. OptoTAF15 droplets exhibit a nearly 4-fold preferential partition of CTD, which is higher than other optoIDRs. The lines indicate the mean and error bars are standard deviations. ** denotes p value of unequal variance t-test < 0.01. (**C**) OptoTAF15-10RD has a higher saturation concentration (C*) than optoTAF15 in nucleoplasm. Inhibition of CTD phosphorylation with THZ1 leads to a decrease in the nucleoplasmic saturation concentration of optoTAF15 and optoTAF15-10RD, but no significant (n.s.) change in cytoplasm. (**D**) The relative cytoplasmic-to-nucleoplasmic saturation concentration correlates with nuclear CTD partitioning into optoIDR droplets. The dashed curve is qualitative.

TAF15 exhibits a distribution of charged amino acid residues along its IDR, notably arginine (R), which is known to interact with tyrosine (Y) residues on CTD through cation-π interactions, and negatively charged aspartic acid (D), which electrostatically interacts with lysine (K) on CTD ^30^. To test those interactions, we mutated optoTAF15 (optoTAF15-10RD) to reduce its charge by replacing arginine (R) with glutamine (Q), and aspartic acid (D) with serine (S) (Fig. S1). CTD partitioning is significantly diminished in the charge-reduced optoTAF15-10RD droplets (Fig. 2A and 2B). Moreover, the optoTAF15-10RD construct has a significantly weakened tendency to phase separate, as indicated by its higher saturation concentration in nucleoplasm (Fig. 2C). Taken together, these data suggest that charge-mediated interactions between TAF15 and CTD contribute to TAF15’s preferential condensation in the nucleoplasm.

Pol II CTD is known to undergo cycles of post-translational modifications to regulate transcriptional activity, including phosphorylation ^27^, which alters charge-mediated interactions. Among these regulators is CDK7, a key kinase which negatively regulates the ability of CTD to function as a transcriptional scaffold ^31^. To test whether the phosphorylation state of CTD influences optoTAF15’s propensity to phase separate, we utilized the CDK7 inhibiting drug THZ1 ^32, 33^. We find that THZ1 treatment decreases the nucleoplasmic saturation concentration of optoTAF15, suggesting unphosphorylated CTD more strongly interacts with optoTAF15 to promotes its phase separation. Consistent with this interpretation, no significant change in the optoTAF15 saturation concentration is observed in the cytoplasm, where CTD is absent (Fig. 2C). Moreover, the partition coefficient of CTD into nuclear droplets strongly correlates with the difference between nucleoplasmic and cytoplasmic saturation concentrations (Fig. 2D). Thus, the degree of CTD phosphorylation modulates phase separation of transcriptional IDRs in the nucleus, and this effect correlates with the apparent interaction between CTD and the transcriptional IDRs.

### Pol II CTD clusters impact the nucleation kinetics of transcriptional condensates

We reasoned that this tendency for CTD to enhance phase separation of TAF15 should exhibit strong spatial variability, as Pol II is not evenly distributed throughout the nucleoplasm, but instead is enriched in transcriptionally active regions ^34, 35^. To examine whether optoTAF15 constructs exhibit any pattern to where and when they condense, we performed blue light cycling experiments where TAF15 condensates are induced to repeatedly phase separate, dissolve, and then again phase separate. We found that some of the optoTAF15 condensates reform in the same physical locations, but that these repetitively forming droplets are transient, and after several cycles they no longer form (Fig. 3 A and 3B). This behavior is in marked contrast to control optoDDX4 droplets, which appear at random locations where they were not found in previous cycles (Fig. S3). To test whether stabilizing endogenous Pol II clusters would alter optoTAF15 behavior in cycling experiments, we treated cells with actinomycin D (Act D), a drug that blocks transcriptional elongation and thus locks Pol II at its promoter-proximal pausing stage ^36, 37^. Act D-treated cells show repetitive localization of optoTAF15 to the same locations in subsequent light illumination cycles, consistent with Pol II remaining locked at high concentrations around specific transcriptional loci, thereby acting as stable nucleation sites for optoTAF15 condensation (Fig. 3C and 3D).

**Fig. 3.**
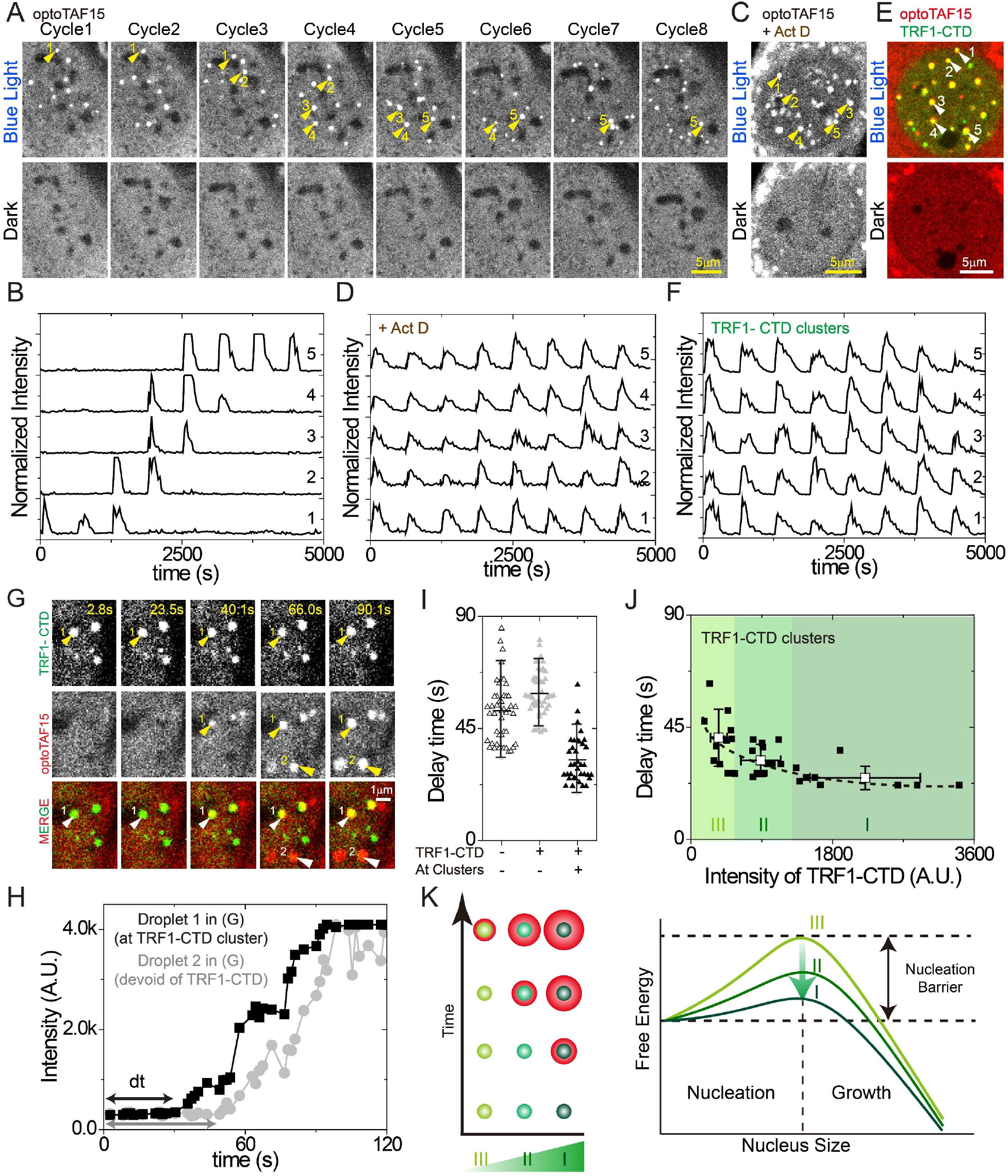
3 Pol II CTD clusters regulate nucleation kinetics of transcriptional condensates. **(A)** Fluorescent images of cells expressing optoTAF15 at the end of each “blue light” (or “dark”) session for 8 cycles are shown. Each cycle consists of 1 min of blue-light illumination (activation) and 9 min of dark (deactivation). Arrowheads point to larger droplets which are analyzed in (B). Scale bar, 5μm. **(B)** Time trace of maximal fluorescence intensity in optoTAF15 droplets throughout 8 cycles of blue light on-and-off; analyzed droplets are indicated by arrowheads in (A). For each trace, intensity is normalized to its maximum. **(C)** Fluorescent images of cells treated with Act D, throughout blue light on-and-off cycles as in (A). **(D)** Time trace for condensates in Act D treated cells shown in (C). **(E)** Fluorescent images of cells co-expressing engineered TRF1-CTD, throughout blue light on-and-off cycles as in (A). **(F)** Time trace for condensates in TRF1-CTD cells shown in (E). **(G, H)** TRF1-CTD sites nucleate significantly faster than optoTAF15 condensates that form elsewhere. Time traces in (H) show maximal fluorescence intensity in two induced optoTAF15 droplets indicated in (G): Black, droplet 1 at a TRF1-CTD cluster; Gray, droplet 2 not associated with any TRF1-CTD cluster. **(I)** Nucleation delay time of optoTAF15. The lines indicate the mean and error bars are standard deviations. Open black symbols indicate the assembly of optoTAF15 condensates in non-TRF1-CTD expressing cells. Black and gray symbols indicate the assembly of optoTAF15 condensates in TRF1-CTD-expressing cells, at TRF1-CTD clusters (black symbols) and elsewhere (gray symbols). **(J)** Nucleation delay time as a function of fluorescence intensity of TRF1-CTD clusters. The dashed curve is qualitative. **(K)** Schematic diagram showing the kinetic effect of nucleation and growth rate during phase separation.

To further examine the hypothesis that native Pol II clusters provide high concentration CTD seeds that nucleate TAF15-rich condensates, we used an engineered system that stably localizes CTD to telomeres, by fusing the telomere associated protein TRF1 to EGFP-CTD (TRF1-CTD). As expected, TRF1-CTD appears as bright puncta, localized at telomeres. In light cycling experiments, the optoTAF15 condensates induced at telomeres always assemble and dissolve at the same locations; unlike the dynamic endogenous Pol II clusters, these engineered CTD foci provide long-term stable nucleation sites for optoTAF15 (Fig. 3E and 3F).

Classical nucleation theory describes how localized “seeds” can decrease the energetic barrier to nucleation, and thus increase nucleation rate (or decrease the delay time until nucleation onset) ^38^. To quantitatively test this prediction of nucleation kinetics, we measured the nucleation delay time (dt), and find that optoTAF15 at TRF1-CTD clusters nucleates significantly faster than optoTAF15 condensates that form elsewhere in the nucleus (Fig. 3G and 3H). The delay time of optoTAF15 condensates formed in these cells away from the TRF1-CTD clusters is similar to that of condensates forming in non-TRF1-CTD expressing cells (Fig. 3 I), dt= 59.9 ± 9.2, and dt= 52.1 ± 13.0 second, respectively. By contrast, the delay time distribution around TRF-CTD clusters are shifted to significantly lower times, dt= 32.4 ± 9.4 second. Moreover, consistent with classical nucleation theory, the delay time is inversely correlated to the concentration of TRF1-CTD clusters, with the brightest clusters nucleating optoTAF15 droplets in 24.7 ± 4.7 second (Group I, Fig. 3J); the dimmest TRF1-CTD clusters nucleate optoTAF15 droplets in 40.8 ± 11.4 second (Group III, Fig. 3J), approaching the long delay times measured for non TRF1-CTD seeding. These results provide further quantitative evidence that clusters of Pol II can modulate and localize the kinetic nucleation of TAF15 condensation (Fig. 3K).

### Condensation of FET-family IDRs enhance transcription

TAF15 is a transcriptional activator, and we wondered whether condensation of optoTAF15 droplets and associated partition of additional of Pol II would impact transcriptional activity. To test this hypothesis, we used 5-ethynyl uridine (EU, a uridine analog) to label nascent RNA transcripts and quantify their spatial location and amount. In cells expressing optoTAF15, but kept in the dark, few nuclei exhibited detectable EU puncta, but instead the EU signal was distributed evenly throughout the nucleoplasm (Fig. S4). By contrast, in blue light-illuminated optoTAF15 cells, an enriched EU signal was detected within optoTAF15 condensates (Fig. 4A and 4B). This enhanced EU signal is due to nascent transcription and is not an artifact of the optoIDR constructs, as no localized EU puncta were detected in cells treated with Act D to inhibit transcription elongation (Fig. S4). Moreover, no puncta were observed in blue light-induced optoDDX4 droplets (Fig. 4A and 4B). OptoEWS and optoFUS showed only a weak EU signal (Fig. 4A and 4B), consistent with our measurements of their weaker CTD recruitment (Fig. 2B). Remarkably, this ability of optoTAF15 condensates to enhance local transcription was also observed in telomere-tethered EGFP-CTD cells (Fig. S4); while telomeres are generally considered transcriptionally quiescent, telomeric repeat-containing RNA (TERRA) is known to be transcribed at telomeres ^39^, and this enhanced EU signal at telomeres could reflect local enhancement of TERRA transcription (Fig. S4).

**Fig. 4.**
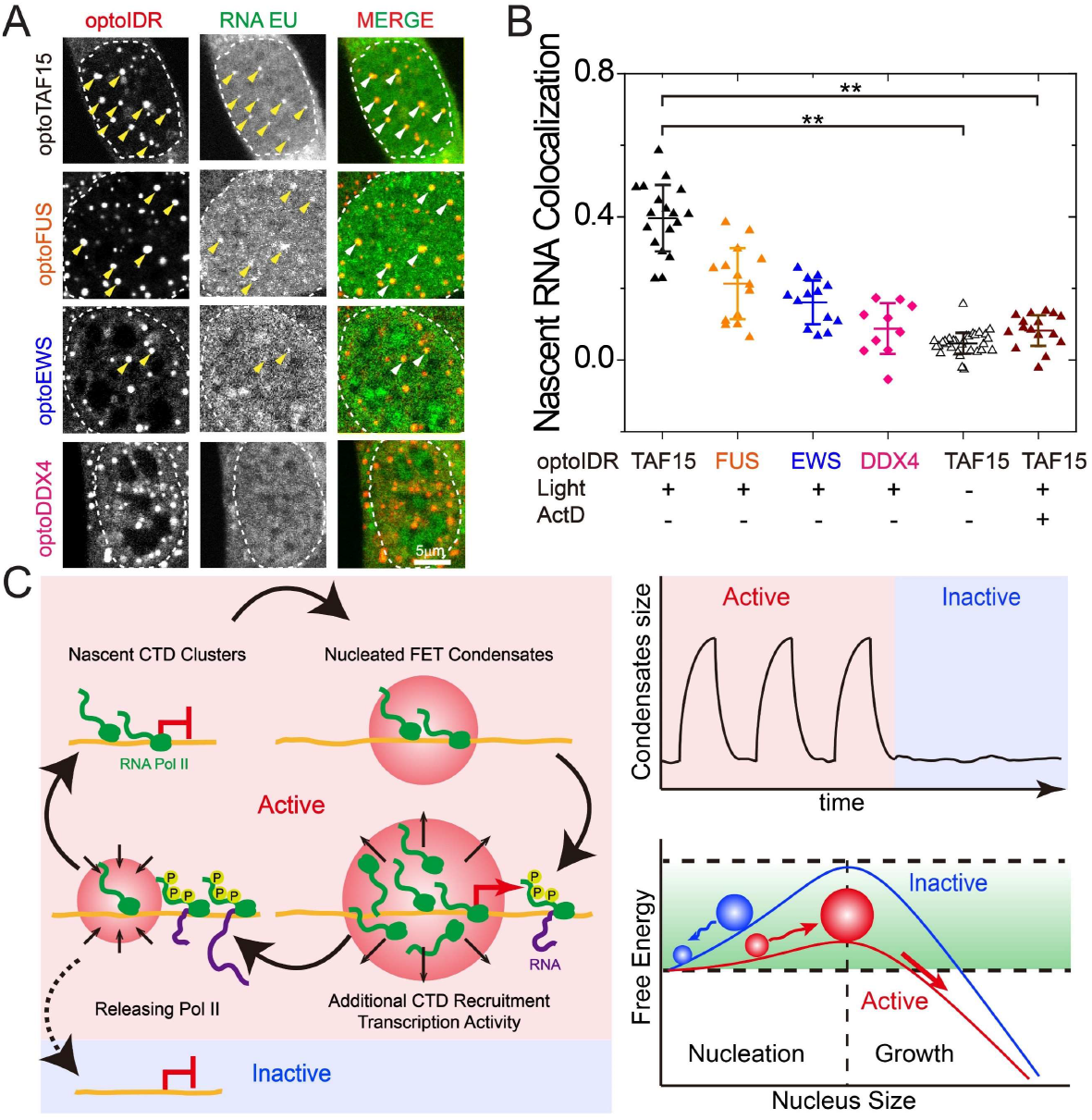
Condensation of transcriptional IDRs enhance transcription. **(A)** Fluorescent images of cell expressing different optoIDR constructs after blue light-illumination. Nascent RNA transcripts are labeled by EU incorporation. Arrowheads point to puncta of nascent transcripts colocalized with optoTAF15 condensates. Cell nucleus is outlined by dotted line. Scale bar, 5 μm. **(B)** Colocalization for nascent RNA various optoIDRs. Degree of colocalization is measured by Pearson’s correlation coefficient of nuclear pixel intensity (nucleoli excluded) between RNA EU and optoIDR channels; +1 indicates perfect correlation, 0 no correlation, and −1 perfect anti-correlation. The lines within indicate the mean and error bars are standard deviations. Significance is denoted by ** for p value of unequal variance t-test < 0.01. **(C)** Conceptual model for the role of transcription condensate nucleation underlying bursts of gene expression. Transcription condensates heterogeneously nucleate at nascent CTD hubs and recruit additional CTD. These amplified transcription loci further drive transcription elongation.

## Discussion

In a number of different experimental systems, transcriptional activity has been found to exhibit intermittent activation ^24, 40^, with bursts of transcription followed by periods of quiescence. We propose that these intermittent dynamics arise from the “all-or-none” nature of nucleated phase separation. Indeed, our data shows that even in nuclei supersaturated with TAF15, phase separation at a particular genomic location only occurs if the nucleation barrier is overcome; however, OptoTAF15 condensates undergo repeated nucleation at the same location for a limited duration, suggesting that the seeding CTD clusters exist only temporarily. Thus, initial accumulation of CTD at a locus can act as a seed for TAF15 condensate nucleation, subsequently driving additional CTD recruitment. Upon transcriptional elongation and native regulation by CDK7, the CTD of endogenous polymerase is phosphorylated and can no longer interact strongly with TAF15. Ultimately, the balance tips, phosphorylation and dispersal of CTDs results in lowered recruitment of TAF15, the condensate dissolves, and the transcriptional burst stops. Upon recruitment of a new Pol II CTD seed, TAF15 and other transcriptional condensates again can be nucleated to repeat the amplification and transcriptional bursting cycle (Fig. 4C).

Our study to quantify intracellular phase diagrams and nucleation kinetics of transcriptional condensates is only the beginning of the union of fundamental biophysics and transcriptional regulation. Future work will move towards precise understanding of the endogenous energetic barrier that must be overcome to initiate transcriptional bursts at single loci, and will model the interplay with non-equilibrium driving forces impacting global transcriptional dynamics.

## Acknowledgements

We thank Michael Levine, Mikko Haataja, Amy Strom and other members of the Brangwynne laboratory for helpful discussions and comments on this manuscript. This work was supported by the Howard Hughes Medical Institute, and grants from the NIH 4D Nucleome Program (U01 DA040601), the Princeton Center for Complex Materials, an NSF supported MRSEC (DMR 1420541), as well an NSF CAREER award (1253035).

## Materials and Methods

### Mammalian cell culture

NIH 3T3 and Lenti-X 293T cells were cultured in growth medium (DMEM, GIBCO) supplemented with 10% FBS (Atlanta Biological) and penicillin and streptomycin (GIBCO) at 37°C with 5% CO_2_ in a humidified incubator.

### Plasmid construction

DNA fragments encoding human TAF15_N_ (IDR of TAF15, residues 1-208), human TRF1 (both from our previous study (*1*)), CTD of the largest subunit of human RNA polymerase II (RPB1, Addgene 35175) and EGFP (Addgene 122147) was amplified by PCR using Phusion High-Fidelity DNA Polymerase (NEB). DNA fragments for the IDR of EWS (residues 47-266) and TAF15_N_-10RD were synthesized (Integrated DNA Technologies). Plasmids for OptoFUS containing IDR of human FUS (residues 1-214) and optoDDX4 containing IDR of DDX4 (residues 1-236) were generated in our previous study (2) and are available through Addgene (101223 and 101225). The sequences of IDRs used in this study are shown in Fig. S1.

All constructs were cloned into pHR-based vector. For optoIDR constructs (IDR-fusion Cry2 plasmids), DNA fragments encoding the IDRs were inserted into the linearized pHR-mCh-Cry2WT (from our previous study (*2*), Addgene 101221) cut with MluI-HF (NEB) and treated with Quick CIP (NEB). All DNA constructs were assembled using In-Fusion HD cloning kit (Clonetech) following manufacturer’s instruction. Assembled products were transformed into Stellar competent cells (Clontech), from which single colonies were picked, grown in LB growth medium supplemented with Ampicillin overnight, and isolated (QIAprep spin miniprep kit, QIAGEN) following manufacturer instructions. All cloning products were confirmed by Sanger sequencing (GENEWIZ).

### Construction of stable cell lines

To produce stable cell lines, lentivirus was produced by transfecting the transfer plasmids, pCMV-dR8.91, and pMD2.G (9:8:1, mass ratio) into Lenti-X 293T cells grown to approximately 70% confluency in 6-well plates. A total of 3 μg plasmid and 9 μL of FuGENE HD Transfection Reagent (Promega) were delivered into each well following manufacturer’s instruction. After 2 days, supernatant containing viral particles was harvested and filtered with 0.45 μm filter (VWR). NIH 3T3 cells plated at ~ 30% confluency in the 12-well plates were infected by adding 1 mL of filtered viral supernatant directly to the cell medium. Virus-containing medium was replaced with fresh growth medium 24 hours post-infection.

### Sample preparation for live cell imaging

Cells were plated on 35-mm glass-bottom dishes pre-coated with Poly-D-Lysine (MatTek) and grown typically overnight in normal growth medium to reach ~50% confluency.

For transcription inhibition experiments, Actinomycin D (Sigma-Aldrich) was dissolved in DMSO at 0.5 mg/mL and further diluted in growth medium to a final concentration of 5 μg/ml before being applied to NIH 3T3 cells followed by a 2-hour incubation prior to being imaged. For CDK7 inhibition experiments, THZ1 (Med Chem Express) was dissolved in DMSO at 1 mM and further diluted with growth medium to a final concentration of 1 μM before being applied to NIH 3T3 cells followed by a 1.5-hour incubation prior to being imaged.

### Nascent RNA transcripts labeling of cultured cells

To label nascent RNA transcripts in live cells, Click-iT RNA Alexa Fluor 488 Imaging Kit was used following protocol from the manufacturer with Alexa Fluor 647 azide from Click-iT EdU Alexa Fluor 647 Imaging Kit. Cells seeded in a 35-mm glass bottom dish were first illuminated by blue light for 15 minutes before 5-ethyl uridine (EU) was added and incubated for 15mins under blue-light illumination. 8 μL of 100 mM EU stock solution was added into each dish containing 2 mL growth medium to obtain a final concentration of 0.4 μΜ. After EU labeling, cells were first washed with PBS, and then fixed and permeabilized using 3.7% formaldehyde and 0.5% Triton X-100 in PBS, respectively, for 15 minutes each at room temperature, and washed again with PBS. Next, Alexa Fluor 647 conjugation was carried out by adding 500 μL of Click-iT reaction cocktail containing Alexa Fluor 647 azide to each dish; samples were then incubated for 30 minutes at room temperature without blue light before being washed with Click-iT reaction rinse buffer and PBS. Finally, DNA was stained with Hoechst 33342 (10 μg/mL in PBS) for 15 minutes at room temperature and samples were washed twice with PBS prior to imaging.

### RNA fluorescence in situ hybridization

RNA fluorescence in situ hybridization (FISH) experiments were performed following protocols from Biosearch Technologies with some modifications. Cells were incubated at 37 °C for 30 minutes with or without blue light illumination and fixed with 4% paraformaldyhyde in PBS at room temperature for 10 minutes. Cells were then permeabilized with 70% ethanol at room temperature for 1 hour and pre-equilibrated with Buffer A (0.2X Stellaris RNA FISH Wash Buffer A and 10% formamide in PBS) at room temperature for 5 minutes. To probe TERRA RNA, hybridization was carried out using 0.125μM DNA probe of the following sequence: 5’-TAACCCTAACCCTAACCCTAACCCTAACCCTAACCCTAACCC-/3Cy5Sp/-3’ (/3Cy5Sp/indicates 3’-end Cy5 modification) in Stellaris RNA FISH Hybridization Buffer with 10% formamide in a humidified chamber incubated at 37°C overnight. Samples were then washed twice with Buffer A at 37°C for 30 minutes; 5 μg/mL DAPI was included during the second wash. Then cells were washed with Stellaris RNA FISH Wash Buffer B at room temperature for 5 minutes. Finally, anti-fade reagent Vectashield was added and the samples were covered by cover-glass and sealed with nail polish. Samples were stored in 4°C for one day before being imaged.

### Microscopy

All images were taken using 60X immersion objective (Plan Apo 60X/1.4, Nikon) on a Nikon A1 laser scanning confocal microscope. An imaging chamber was maintained at 37°C with 5% CO_2_. For activation, cells were usually imaged with 440-nm laser with a dichroic filter for the 488-nm laser in order to attenuate blue laser intensity at the specimen plane below 0.4 μW.

For studying light-activated phase behavior, the activation protocol was performed with 440-nm laser every second and fluorescent image of mCherry was acquired with 568-nm laser every 10 seconds, with pin hole size set to 33.2 μm, frame size to 512×512 pixel-square, and dwell time to 2.2 μs. For each cycle of a multi-cycle activation, the above activation protocol was applied for 1 minute, followed by a 9-minute deactivation (acquisition of mCherry fluorescent image every 10 seconds); this cycle was repeated for 8 times.

For delay time of initial nuclearization measurement, activation protocols were performed with 440 nm per each 10 second and with 568 nm per each second to observe mCherry channel with pin hole size set to 33.2 μm, 512×512 pix frame sizes, and 2.2 μs dwell time.

To observe multi-channel image, 667-nm, 568-nm, 488-nm, and 405-nm lasers were used to excite Alexa Fluor 647, mCherry, EGFP, and DAPI (or Hoechst 33342) in cells, respectively.

### Fluorescence Correlation Spectroscopy

We used a laser scanning confocal microscope (Nikon A1; an oil immersion objective, Plan Apo 60X/1.4, Nikon) combined with Fluorescence Correlation Spectroscopy (FCS) Upgrade Kit for Laser Scanning Microscopes (PicoQuant). FCS measurements were performed using the SymPhoTime Software (PicoQuant). Data for diffusion and concentration of proteins were obtained with 30 s of FCS measurement time. For each measurement, the resulting auto-correlation functions were fitted with the pure diffusion model. The measurement volume was determined by fluorophore dye Atto 550 in water (ω_xy_ = 0.18 μm and the ratio of axial to radial of measurement volume, κ = ω_z_/ω_xy_ = 5.1). Here, in living cells, the refractive index mismatch that can distort the FCS measurement volume as a non-Gaussian profile as well as other optical artifacts would lead to an absolute error of ~20% in diffusion coefficient and concentration(*3*).

The dependence of the diffusivity (D) on protein concentration (C) can be written as: D = D_0_ (1 + βC), where D is protein diffusivity, D_0_ is the diffusivity at infinite dilution, β is interaction strength, and C is the protein concentration. β can be written as: (2B_2_M − ks − υ) Mc, where M is the molecular weight, B_2_ is the second virial coefficient, ks is sedimentation interaction parameter, υ is partial specific volume, and c is protein weight concentration (*4*). Mean and standard deviation of diffusion coefficient and concentration were inferred from fitting correlation curves from 5-10 different cells. β was calculated by taking the slope of the fitted lines of diffusion coefficient vs. concentration for each cell line.

### Image analysis of molecular concentrations

Total concentrations of molecules as well as steady-state background concentrations outside clusters were measured from fluorescence images of cells using ImageJ (NIH), and corrected by subtracting background noises measured with areas devoid of any cells. Cells expressing optoIDRs rapidly assemble optoDroplets under blue light illumination as shows in Fig. S2.

### Colocalization analysis

The colocalization coefficient between two-color images was estimated by calculating Pearson’s correlation coefficient (PCC) defined as;

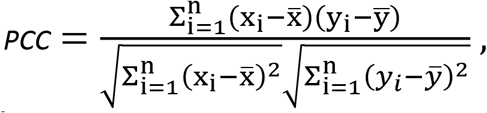

where x_i_ and 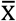 are the i^th^ pixel intensity and mean values in the segmented color 1 image, respectively. Likewise, y_i_ and 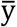 are the i^th^ pixel intensity and mean values in the color 2 image, respectively. The perfect-correlation, no-correlation, and anti-correlation give the values of PCC = 1, 0, −1, respectively. The correlation in the whole nucleus (except for nucleoli) was calculated (*5*).

---

**Fig. S1.**
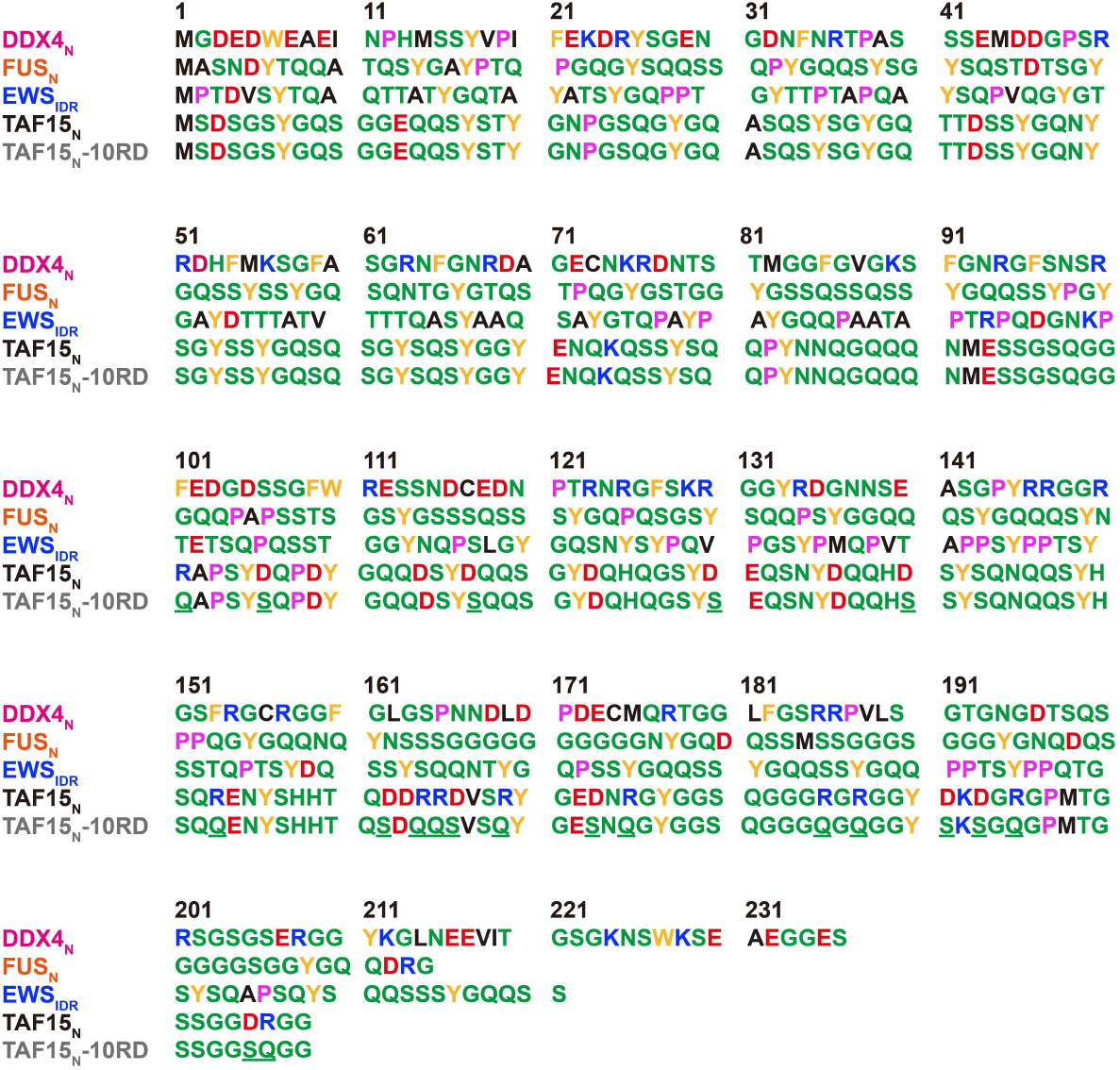
The amino acid sequence of intrinsically disordered regions. Polar residues are shown in green; positively charged are shown in blue; negatively charged are shown in red; hydrophobic are shown in black; aromatic residues are shown in orange.

**Fig. S2.**
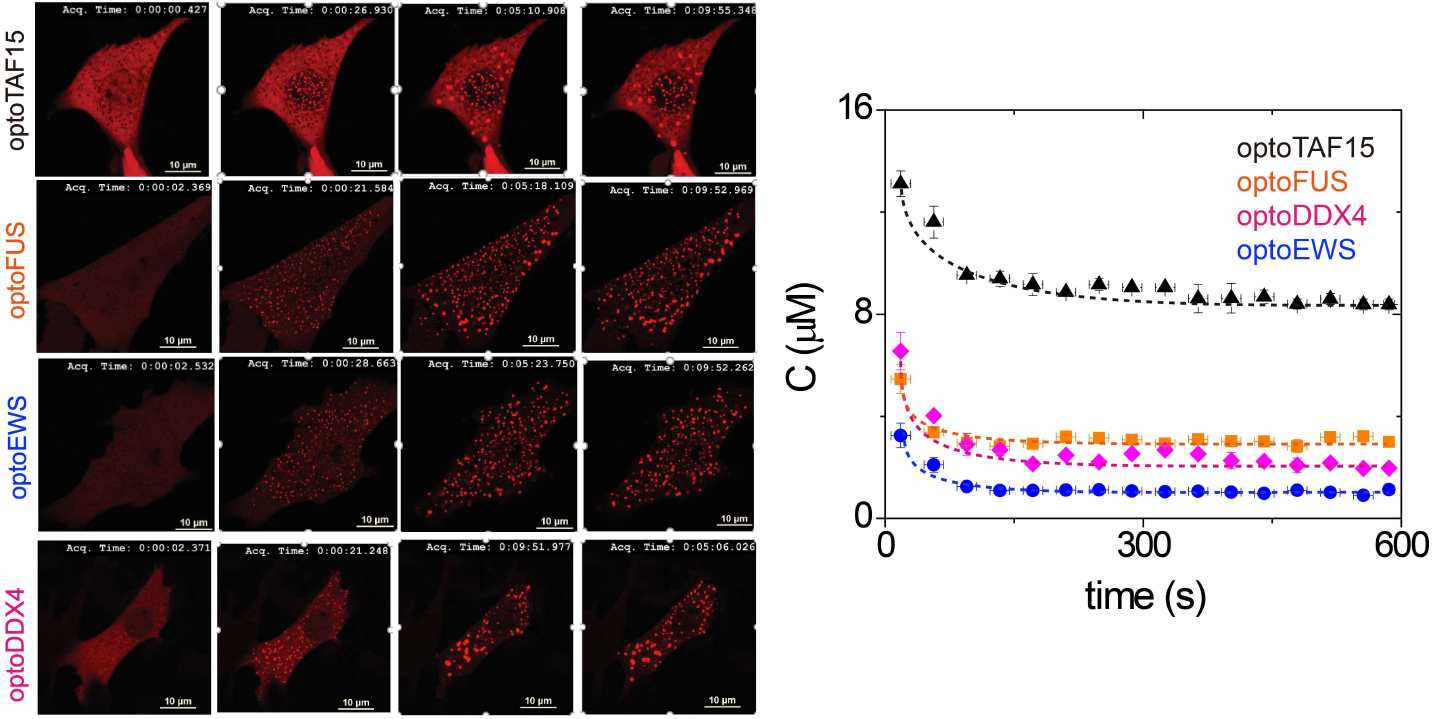
Rapid clustering of optoIDRs upon blue light illumination. Images of optoIDRs cells exposed by blue light illumination. Time since the start of illumination is shown with 0 sec indicating before illumination. The protein concentration was determined by the fluorescence intensity outside of clusters. The concentration was measured during clustering until it reaches steady state.

**Fig. S3.**
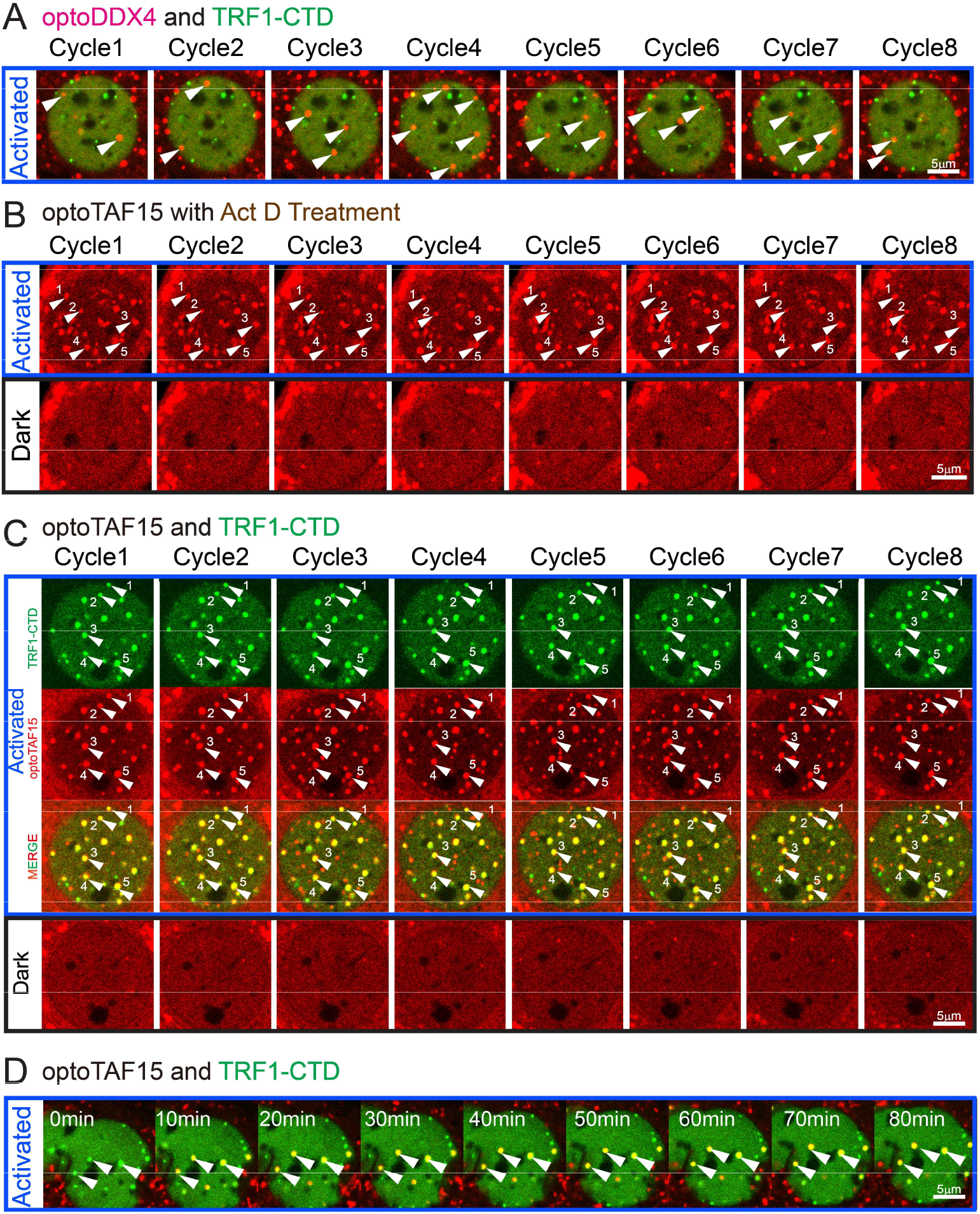
Repeated assemble and disassemble of optoDroplets. **(A)** OptoDDX4 and TRF1-CTD cells, **(B)** Act D treated optoTAF15 cells, and **(C)** optoTAF15 and TRF1-CTD cells were exposed to blue light activation condition for 1 min to assemble clusters and then incubated in the absence of blue light for 9 min to disassemble clusters. Cell images before and after activation at the end of each cycle are shown. **(D)** OptoTAF15 and TRF1-CTD cells were exposed to continuous blue light illumination. OptoTAF15 condensates remain stable at telomere tethered CTD sites. Scale bars, 5μm.

**Fig. S4.**
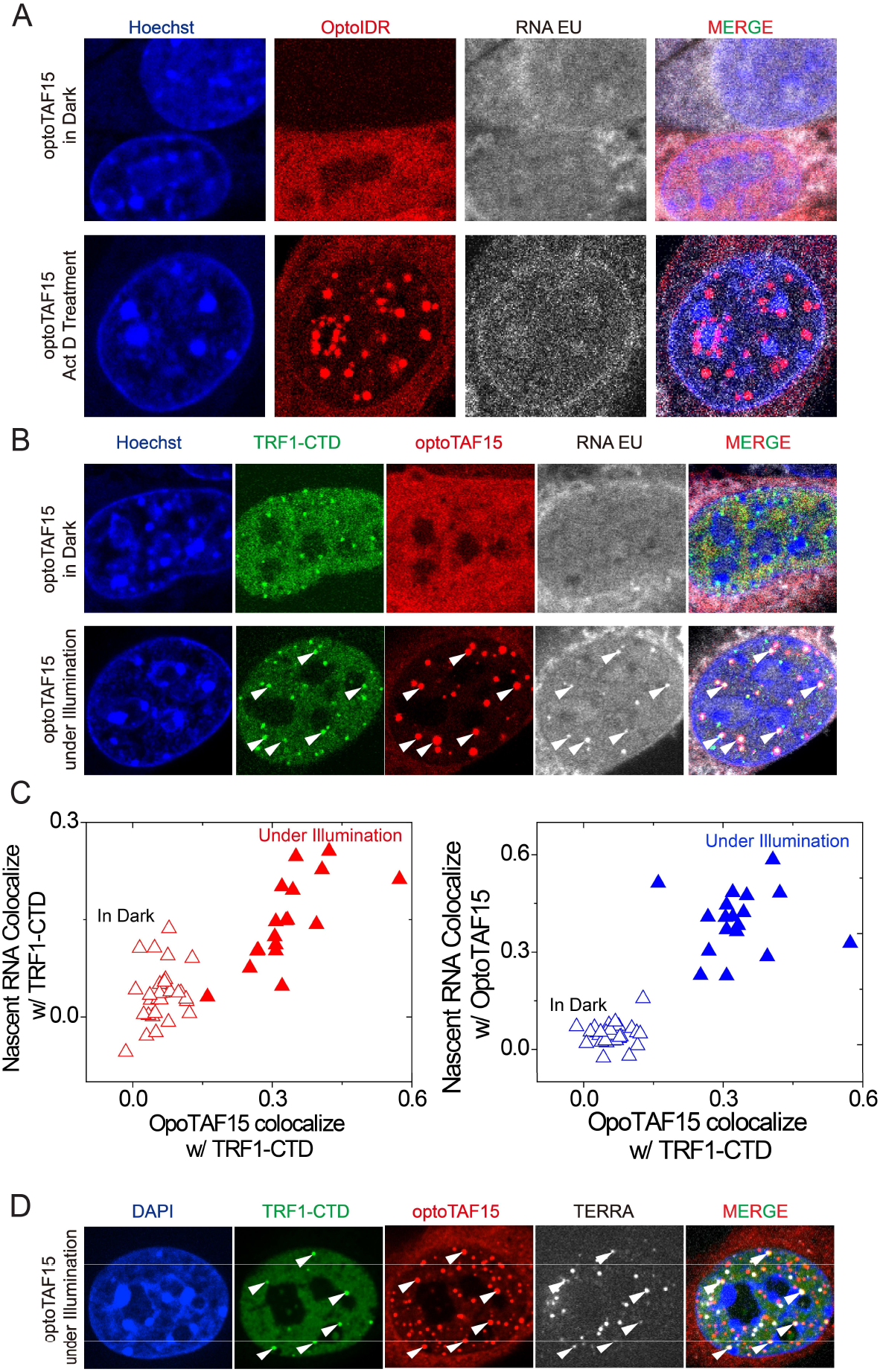
Image of cell with nascent RNA production labeled by EU incorporation. **(A)** In cells expressing optoTAF15 and kept in dark, few of them exhibited detectable localized puncta of EU, but instead the EU signal was distributed evenly throughout the nucleoplasm. In Act D treated cells under blue light illumination, nascent RNA transcripts were not present in optoTAF15 condensates. **(B)** In cells expressing optoTAF15 and TRF1-CTD and kept in dark, the EU signal was distributed evenly throughout the nucleoplasm. Image of cell with nascent RNA production labeled by EU incorporation. In cells expressing optoTAF15 and TRF1-CTD under blue light illumination, nascent RNA transcripts were present in optoTAF15 condensates. White arrows indicate light-illuminated TAF15 condensates colocalize TRF1-CTD clusters and enhance local transcription RNA EU clusters. **(C)** Colocalization results indicate that light-illuminated optoTAF15 condensates strongly colocalize with local nascent RNA EU production at TRF1-CTD clusters. **(D)** White arrows indicate light-illuminated TAF15 condensates colocalize TRF1-CTD clusters and enhance local transcription TERRA at telomeres.

